# Sepsis Increases Perioperative Metastases in a Murine Model

**DOI:** 10.1101/142083

**Authors:** Lee-Hwa Tai, Abhirami A. Ananth, Rashmi Seth, Almohanad Alkayyal, Jiqing Zhang, Christiano Tanese de Souza, Phillip Staibano, Michael A. Kennedy, Rebecca C. Auer

## Abstract

**Background:** Cancer surgery can promote tumour metastases and worsen prognosis, but the effects of perioperative complications such as sepsis, blood loss, and hypothermia on subsequent cancer metastases have not been addressed.

**Objective:** To evaluate the effect of common perioperative factors on postoperative tumour metastases in murine models of cancer surgery. We hypothesize that perioperative blood loss, hypothermia, and sepsis facilitate tumour metastases in these murine models.

**Methods:** Prior to surgery, pulmonary metastases were established by intravenous challenge of CT26LacZ colon cancer cells in Balb/c mice or B16LacZ melanoma cells in C57Bl/6 mice. Surgical stress was generated through partial hepatectomy (PH) or left nephrectomy (LN). Sepsis was induced by puncturing the cecum to express stool into the abdomen. Hemorrhagic shock was induced by removal of 30% of total blood volume via saphenous vein. Hypothermia was induced by removing the heating apparatus during surgery and lowering core body temperatures to 30°C. Lung tumour burden was quantified 3 days following surgery.

**Results:** Surgically stressed mice subjected to Stage 3 hemorrhagic shock or hypothermia did not show an additional increase in lung tumour burden. In contrast, surgically stressed mice subjected to intraoperative sepsis demonstrated an additional 2-fold increase in the number of tumour metastases. Furthermore, natural killer (NK) cell function, as assessed by YAC-1 tumour cell lysis, was significantly attenuated in surgically stressed mice subjected to intraoperative sepsis. Both NK cell-mediated cytotoxic function and lung metastases were improved with perioperative administration of polyI:C, which is a ligand for toll-like receptor (TLR)-3

**Conclusions:** Perioperative sepsis, but not hemorrhagic shock or hypothermia, enhances the prometastatic effect of surgery in murine models of cancer. Identification of the mechanisms underlying perioperative immune suppression will be critical in the development of immunomodulation strategies that aim to attenuate perioperative metastatic disease.

## INTRODUCTION

Severe trauma causes compensatory changes in immune, neural, endocrine, and metabolic function[1]. Similarly, surgical stress can lead to the onset of prothrombotic and immunosuppressive changes during the postoperative period[2, 3]. Many correlative studies have confirmed an association between postoperative complications, immune suppression, and worsened cancer survival[4–7]. Our group and others have proposed surgery-induced cellular immune suppression as a primary factor in the progression of cancer, including local recurrence and metastases[8–20]. In humans, suppression of the cellular immune response following major surgery appears to peak at 3 days[21], but can also persist for weeks[17, 22, 23]. These immunosuppressive changes are characterized by an imbalance in plasma cytokine levels (i.e. a decrease in the levels of interleukin (IL)-2[24] and IL-12[25] and an increase in the levels of IL-6[24, 26, 27] and IL-10[28]) and a decrease in the number and function of circulating CD8+ T cells[29], dendritic cells[30], and natural killer (NK) cells[8, 12, 31]. Specifically, our group reported on the association between coagulation and NK cell function in the development of metastases following cancer surgery[8]. More recently, we employed validated murine models of surgical stress and spontaneous metastases[11] to provide *in vivo* evidence of global NK cell dysfunction in postoperative metastatic disease[32].

Modern surgical techniques minimize the potentially adverse consequences of perioperative events such as intraoperative blood loss, sepsis, and hypothermia. Despite this, severe intraoperative blood loss occurs in approximately 6–10% of patients with advanced cancer[33]. Every year, two million operations are complicated by infection in the US and surgery accounts for 30% of sepsis diagnoses[34]. Furthermore, 8.5% of all cancer-related deaths are due to the concurrent onset of severe sepsis[35]. Hypothermia, which is defined as a core body temperature of <36°C, is a common occurrence during surgery, as up to 70% of patients are hypothermic on admission to the recovery room[36].

Clinical studies in cancer patients have confirmed an association between perioperative factors such as hypothermia[37], blood loss[38, 39], and postoperative infections[40, 41], and increased cancer recurrence and reduced cancer-specific survival following cancer surgery.

Despite the epidemiological evidence linking perioperative complications with increased surgical stress and worsened cancer outcomes, the role of intraoperative blood loss, sepsis, and hypothermia in immunosuppression and metastases has not been addressed directly. Our study incorporates three surgical murine models of CRC to investigate the effect of blood loss, sepsis, and hypothermia on NK cell function and metastatic disease. Taking measures to reduce perioperative complications and/or employing preoperative neoadjuvant immunotherapy will help to improve survival outcomes and reduce recurrences of cancer.

## METHODS

### Cell lines

CT26.LacZ mouse colon carcinoma and YAC-1 mouse lymphoma cells were purchased from the American Type Culture Collection (ATCC, Manassas, VA, USA). CT26.LacZ cells were cultured in HyQ high glucose Dulbecco’s modified Eagles medium (GE healthcare, Mississauga, ON, CA) supplemented with 10% fetal bovine serum (CanSera, Etobicoke, ON, CA). YAC-1 cells were cultured in HyClone™ Roswell Park Memorial Institute medium (RPMI)-1640 medium (GE healthcare, Mississauga, ON, CA) supplemented with 10% fetal bovine serum (CanSera, Etobicoke, ON, CA) and 1x of Penicillin/Streptomycin (Invitrogen, Carlsbad, CA, USA).

### Animals

Female age-matched (6–8 weeks old at study initiation) BALB/c mice (Charles River Laboratories, Wilmington, MA, USA) were housed in specific pathogen-free conditions. Animal studies complied with the Canadian Council on Animal Care guidelines and were approved by the University of Ottawa Animal Research Ethics Board.

### Induction of experimental metastasis and surgical stress

Mice were subjected to 2.5% isofluorane (Baxter Corporations, Mississauga, ON, CA) for the induction and maintenance of anesthesia. Routine perioperative care for mice, including the subcutaneous administration of buprenorphine (0.05 mg/kg) for pain control the day of surgery and every 8 hours for 2 days following surgery, was conducted in concordance with University of Ottawa protocols. Surgical stress was induced via an abdominal laparotomy (i.e. 3-cm midline incision), which was preceded by an intravenous challenge with 3e5 CT26.lacZ cells to establish pulmonary metastases. Abdominal laparotomy was commenced 10 minutes following tumor inoculation as previously described[11]. Animals were euthanized at 18 hours or 3 days following tumor inoculation and their lungs were stained with X-gal (Bioshop Canada Inc., Burlington, ON, CA), as described previously[42]. The total number of surface metastases on the largest lung lobe (left lobe) were quantified using a stereomicroscope (Leica Microsystems, Richmond Hill, ON, CA).

### Hypovolemic stress model

Hypovolemic shock was induced by preoperatively bleeding mice prior to tumour inoculation. Mice were bled either 20% (300 uL) or 30% (450 uL) of their total blood volume by puncturing the saphenous vein just above the foot. Systolic arterial pressure (SAP) in conscious mice before and after saphenous vein bleeding was measured using a tail-cuff sphygmomanometer. To do this, mice were kept in a warmed black box and an inflatable cuff was applied to the base of the tail. The tail was then placed on a piezoelectric sensor for analysis of the pressure waveforms.

### Hypothermia stress model

Intraoperative hypothermic shock was induced by placing mice directly on the metal surgical surface without a heating pad immediately following tumour inoculation. Mice were kept under hypothermic conditions and anesthesia for approximately 2 hours and were subsequently housed under normothermic conditions. Rectal temperatures were recorded every 15 minutes throughout the procedure to verify that hypothermia was maintained.

### Sepsis stress model

Intraoperative polymicrobial sepsis was induced in mice by cecal puncture at the time of abdominal laparotomy (i.e. 3-cm midline incision). Polymicrobial sepsis was confirmed by Gram stain of peritoneal lavage fluid, which was isolated 18 hours following surgery. Bacterial counts were determined by serial dilution of peritoneal lavage fluid and overnight culture on tryptic soy broth agar plates at 37°C.

### Ex-vivo NK cell cytotoxicity assay

Chromium release assays were conducted as previously described[43]. Briefly, splenocytes were isolated from surgically stressed and control mice 18 hours after surgery. Pooled and sorted NK cells were resuspended at a concentration of 2.5×10^6^ cells/mL. These cells were then mixed with chromium-labeled YAC-1 target cells, which were resuspended at a concentration of 3×10^4^ cells/mL at various effector-to-target (E:T) ratios (i.e. 50:1, 25:1, 12:1, 6:1).

### Statistical analysis

Statistical tests were performed using GraphPad Prism (GraphPad, San Diego, CA, USA). One-way ANOVAs with Bonferroni multiple comparison tests as well as Student’s t-tests with equal variances were conducted. Data were reported as the mean ± standard error of the mean (SEM). An alpha value of < 0.05 was considered to be statistically significant.

## RESULTS

### Severe hypovolemia increases pulmonary metastases, but is not additive when combined with surgical stress

To discern the effect of hypovolemic shock on perioperative metastases, mice were first exsanguinated through the saphenous vein while systolic blood pressure was measured using a tail cuff sphygmomanometer (Figure 1A). A significant reduction in tail cuff blood pressure was observed following the loss of 450 uL of blood, which is representative of 30% of the total blood volume of a 25 g mouse (71.67 ± 2.186 mmHg vs. 107.7 ± 1.453 mmHg, *p* = 0.0002); moreover, the reduction in tail cuff blood pressure was exacerbated when 450 uL of blood loss was combined with surgical stress (61.67 ± 1.667 mmHg vs. 107.7 ± 1.453 mmHg, *p* <0.0001) (Figure 1B). In the absence of surgical stress, hypovolemic changes due to 30% blood loss significantly increased the number of pulmonary metastases when compared to mice that did not undergo blood loss (438.3 ± 56.89 vs. 205.0 ± 24.88, *p* = 0.0212) (Figure 1C). In addition to hypovolemia, surgical stress alone significantly increased pulmonary metastases compared to mice that did no undergo surgery (287.5 ± 39.01 vs. 95.40 ± 32.35, *p* = 0.0065). However, a combination of 30% blood loss and surgical stress did not further increase the number of pulmonary metastases above surgical stress alone (285.6 ± 35.94 vs. 287.5 ± 39.01, *p* = 0.9725) (Figure 1D).

**Figure 1.**
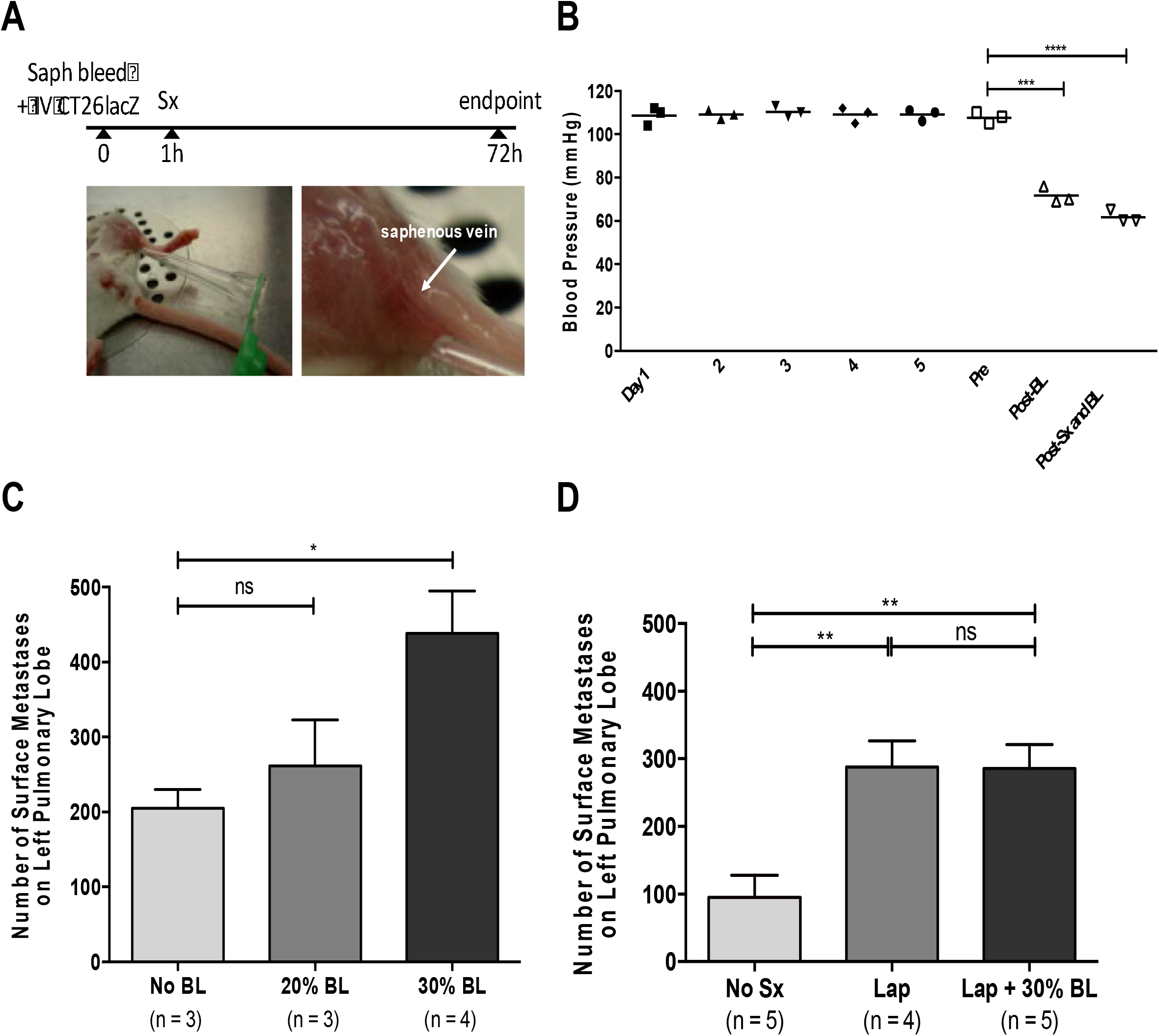
Hemorrhagic shock does not increase metastatic disease. **A. *Experimental overview***. Balb/C mice were bled through the saphenous vein (indicated by the white arrow) and subsequently injected intravenously (IV) through the tail vein with 3x10^5^ CT26lacZ cells. Approximately 1 hour later, surgical stress (sx) was generated by laparotomy (Lap) (5 cm incision). Mice were sacrificed at 72h to quantify lung metastases. **B. *Blood pressure is reduced following surgical stress and blood loss***. Blood pressure (mmHg) was measured following a 5 day training period (Day 1 – 5), prior to bleeding (Pre), immediately following bleeding (Post-BL), and immediately following surgical stress (Post-Sx and BL, n=3). **C. *Blood loss increases metastic burden***. Lung metastases were measured on Day 3 following no blood loss (no BL) or 20% (20% BL) or 30% blood loss (30% BL). **D. *Blood loss does not increase metastatic disease associated with surgical stress***. Lung metastases were measure on Day 3 in mice that did not undergo surgical stress (No Sx) and animals undergoing a laparotomy (Lap) alone or in combination with 30% blood loss (30% BL). Error bars represent ± SEM.

### Severe hypothermia does not increase pulmonary metastases

Next, we sought to assess the effect of perioperative hypothermic shock on metastases in a surgical murine model. Anaesthetized animals were maintained at either normothermic (35.22°C–37.95°C) or hypothermic (26.35°C–29.03°C) temperatures for a 3-hour period following injection of the tumour cells and surgical stress (Figure 2A and 2B). We observed that hypothermia alone did not significantly increase pulmonary metastases when compared to normothermic mice (27.75 ± 7.667 vs. 21.38 ± 7.720, *p* = 0.5673) (Figure 2C). Surgical stress in normothermic conditions did significantly increase the number of pulmonary metastases compared to mice that did not undergo surgery (391.5 ± 38.18 vs. 136.4±21.47, *p* = 0.0024) (Figure 2D). When combined with surgical stress, however, hypothermia did not further increase the number of pulmonary metastases when compared to mice that underwent surgical stress under normothermic conditions (391.5 ± 38.18 vs. 341.8 ± 80.90, *p* = 0.6265).

**Figure 2.**
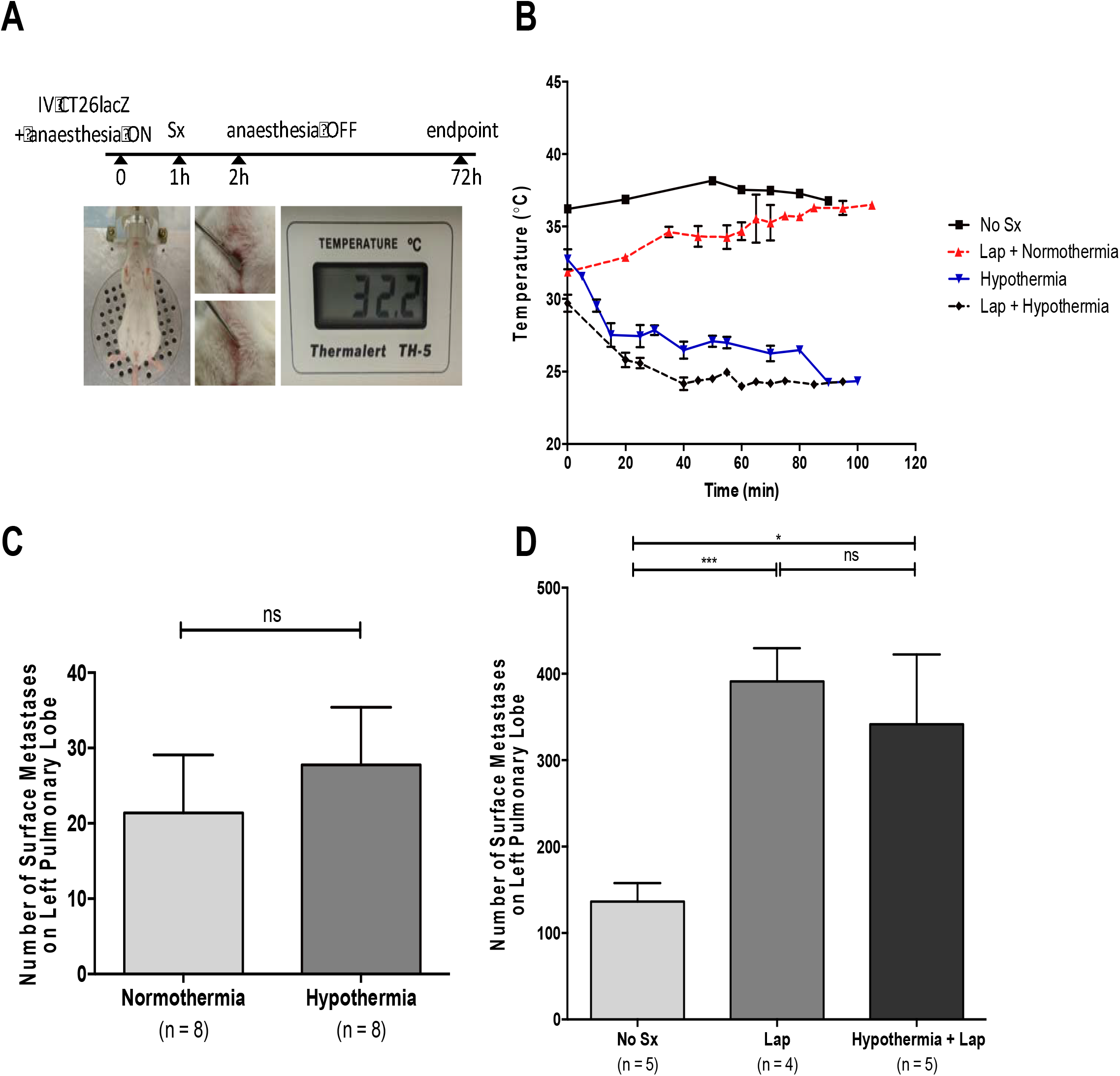
Hypothermia does not increase metastatic disease. **A. *Experimental overview***. Balb/C mice were injected intravenously (IV) through the tail vein with 3x10^5^ CT26lacZ cells. Mice undergoing hypothermia were placed directly on the metal block (shown) while control mice were kept under normothermic conditions with heating pads. One hour into anesthesia, surgical stress (Sx) was generated by laparatomy (Lap). Mice were removed from anesthesia after 2 hours total and kept in normal housing conditions. Temperature was measured during anesthesia using a rectal probe (as shown). Mice were sacrificed at 72h to quantify lung metastases. **B. *Maintenance of body temperature***. Mice kept under normothermic conditions alone ranged from 31.88°C to 38.16°C in comparison to hypothermic conditions which ranged from 23.98°C to 32.75°C for 100 min. **C. *Hypothermia alone does not increase lung metastases***. The impact of body temperature upon lung metastases was assessed in mice maintained at normothermic and hypothermic conditions. **D. *Prometastic effects of surgical stress are not impacted by hypothermic conditions***. Lung metastases were measured in mice that did not undergo surgery (No Sx) and mice undergoing a laparotomy under normothermic (Lap) and hypothermic conditions (Hypothermic + Lap). Error bars represent ± SEM.

### Perioperative polymicrobial sepsis increases pulmonary metastases compared to surgical stress alone

Next, we assessed the effects of sepsis on pulmonary metastases by contaminating the peritoneal cavity with stool expressed through caecal puncture (Figure 3A). Peritoneal lavage contained both Gram-negative and Gram-positive bacteria, as well as coccus and bacillus-shaped bacteria, confirming that the caecal puncture lead to polymicrobial sepsis (Figure 3B). Surgical stress alone resulted in a significant increase in pulmonary metastases when compared to control mice (374.3 ± 20.68 vs. 266.4 ± 16.15, *p* = 0.0006); furthermore, we observed that a combination of polymicrobial sepsis and surgical stress resulted in a significant increase in pulmonary metastases when compared to mice that underwent surgical stress in the absence of sepsis (480.3 ± 18.98 vs. 374.3 ± 20.68, *p* = 0.0010) (Figure 3C). Previous studies have shown that suppression of NK cell cytotoxic activity is responsible for the increase in cancer burden following surgical stress [30]. In this study, we demonstrate that sepsis, in conjunction with surgical stress, significantly attenuated NK cell cytotoxic activity below that of surgical stress alone (Figure 3D).

**Figure 3.**
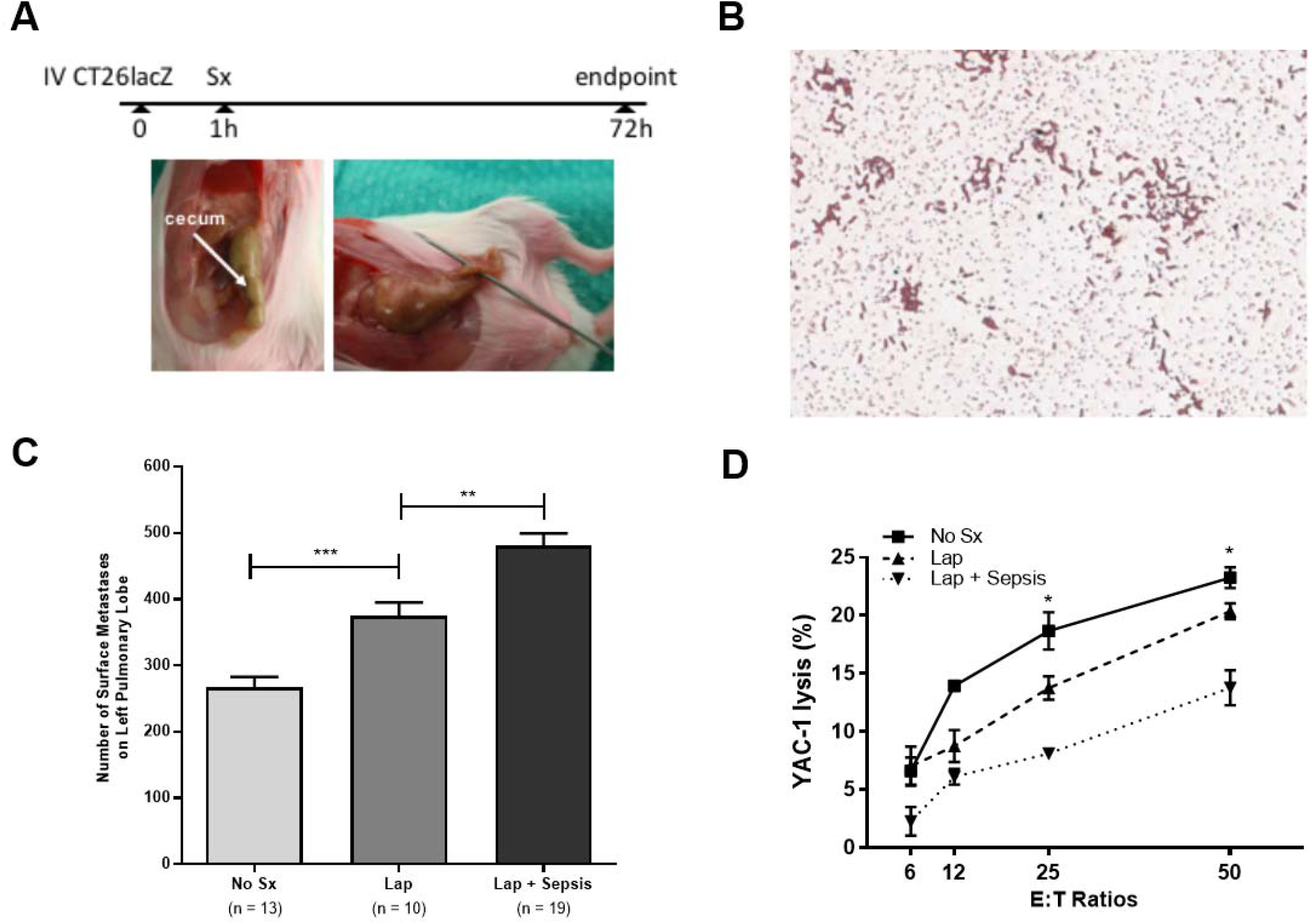
Sepsis increases metastatic disease burden. **A. *Experimental overview***. Balb/C mice were injected intravenously (IV) through the tail vein with 3x10^5^ CT26lacZ cells. One hour later, surgical stress (Sx) was generated by laparatomy (Lap) or laparatomy with caecal puncture (Lap+CP). The caecum was externalized and punctured using an 18G needle (as shown). Mice were sacrificed at 72h to quantify lung metastases. **B. *Detection of bacteria in the peritoneal cavity***. A representative Gram stain of peritoneal lavage fluid from a mouse that underwent Lap+CP indicates polymicrobial sepsis due to the presence of both cocci and bacilli. **C. *Sepsis increases lung metastases***. Lung metastases were quantified in mice that did not undergo surgical stress (No Sx), laparotomy alone (Lap) or laparotomy combined with sepsis (Lap + Sepsis). D. ***Intraoperative sepsis impairs NK cell-mediated cytotoxicity***. Chromium release killing assay of YAC-1 tumour target cells by DX5+ NK cells isolated from animals that did not undergo surgical stress (No Sx), laparotomy alone (Lap) in the presence or absence of sepsis (Lap + Sepsis). Effector (DX5+ NK cells) to target (Yac1) (E:T) ratios of 6:1, 25:1, and 50:1 are shown. Error bars represent ± SEM.

### Perioperative NK cell stimulation reduces metastases and restores NK cell function in the presence of sepsis

We have shown that postoperative immune dysfunction can be ameliorated through the use of NK cell-stimulating agents such as polyI:C, a double-stranded RNA mimetic. In this study, we examined whether sepsis-induced postoperative pulmonary metastases could be suppressed by perioperative immune stimulation with polyI:C (Figure 4A). We demonstrated that treatment with polyI:C alone significantly decreased the number of pulmonary metastases compared to mice undergoing surgical stress and sepsis with no perioperative therapy (18.25 ± 8.390 vs. 175.0 ± 13.48, *p* = 0.0002); moreover, antibiotics alone did not affect the number of pulmonary metastases when compared to surgically stressed mice with sepsis that were not administered perioperative therapy (185.3 ± 19.38 vs. 175.0 ± 13.48, *p* = 0.6794) (Figure 4A). Furthermore, antibiotic therapy did not further decrease the number of pulmonary metastases in mice undergoing surgical stress and sepsis in the presence of polyI:C (25.80 ± 4.306 vs. 18.25 ± 8.390, *p* = 0.4213). In support of a role for NK cells in mediating this effect we also demonstrated that the addition of polyI:C enhanced NK cell cytotoxic activity by 10-fold, a magnitude similar to the reduction in metastases seen with perioperative use (Figure 4B).

**Figure 4.**
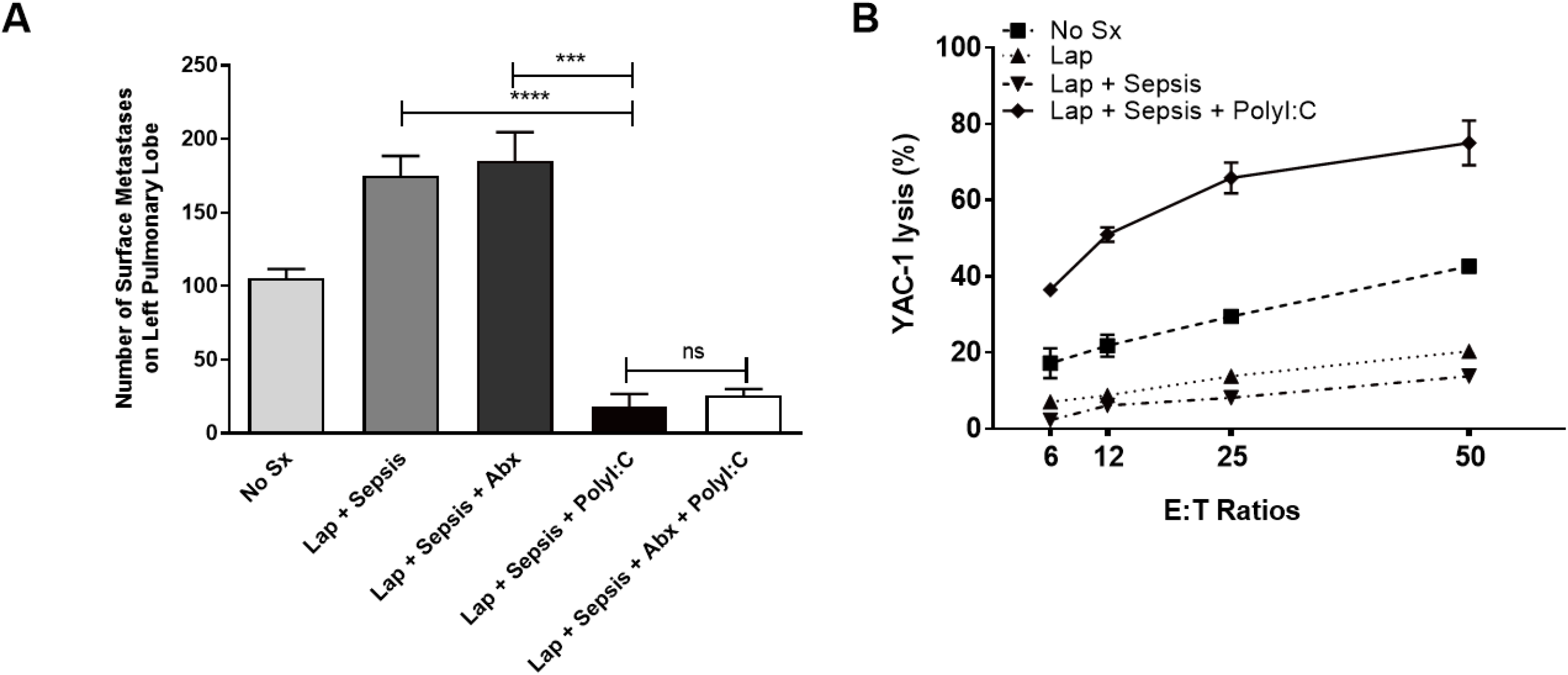
Prometastatic effects of sepsis are reversed with perioperative NK cell activation. **A. *Lung metastases are reduced in polyI:C treated animals***. Lung metastases were quantified in untreated animals (No Sx) and following laparotomy (Lap) without or with perioperative antibiotic treatment administered subcutaneously (Abx, Imipenem 0.5 mg) and preoperative polyI:C treatment (PolyI:C,150 μg/200 μL PBS). **B. *Preoperative PolyI:C restores NK cell cytotoxicity***. Chromium release killing assay of YAC-1 tumour target cells by DX5+ NK cells isolated from animals undergoing treatments as above. Effector (DX5+ NK cells) to target (Yac1) (E:T) ratios of 6:1, 12:1, 25:1, and 50:1 are shown. Error bars represent ± SEM.

## DISCUSSION

Perioperative complications, specifically infection, decrease long-term survival[5, 44] and promote tumour[45] recurrence in patients with CRC. Although hypovolemia in the absence of surgical stress did lead to an increase in pulmonary metastases, our findings demonstrate that neither severe intraoperative hypovolemia nor hypothermia impact the prometastatic effects of surgical stress. Correlative clinical studies confirm that postoperative infections following surgery can accelerate the time to cancer recurrence[46–48]. Here, using murine models we demonstrate that polymicrobial sepsis in conjunction with surgical stress facilitates the development of perioperative lung metastases. Our results suggest that the combined immunosuppressive effects of surgical trauma and sepsis dampen anti-tumour immune responses, ultimately leading to an increase in metastases. In addition to the immunosuppressive effects of surgical stress, severe sepsis can induce lymphocyte exhaustion[49], apoptosis of immune cells[50, 51], and a predominance of immunoregulatory cells, including regulatory T cells[52, 53] and myeloid-derived suppressor cells[54]. This highly suppressive environment likely worsens the already immunosuppressive environment present in most cancer patients in need of surgical intervention[55, 56]. Thus, the immunosuppressive effects of surgery, sepsis, and cancer may interact to severely dampen immune activation and increase the likelihood of cancer recurrence and metastatic disease.

Our findings also suggest that sepsis induces its prometastatic effect by inhibiting NK cell cytotoxic function. In the cancer microenvironment, the anti-tumour function of NK cells is suppressed[57], while a decrease in NK cell number and function in patients undergoing surgery for CRC is associated with heightened mortality and cancer recurrence suggesting that the suppressive effects of sepsis likely exacerbate the already impaired NK cell function[58, 59]. In agreement with our findings, previous studies have demonstrated that sepsis in a non-surgical context can impair NK cell cytotoxicity[60], a finding that has been attributed to a heightened activation of regulatory cell subsets[61]. In particular, murine sepsis models have shown that an increase in regulatory T cells contributes to post-sepsis immunosuppression and potentiates tumour growth[62]. NK cells are a critical component of anti-tumour immunity and so, based on our findings, we suggest that the inhibition of NK cell function is a key player in perioperative cancer recurrence following surgical stress and septic insult.

Tumour-infiltrating NK cells and lymphocytes are associated with improved prognosis several malignancies[63–67]. The enhancement of preoperative NK cell activation with PolyI:C, a TLR3 ligand, to counteract the immunosuppressive effects of surgery and sepsis and attenuate perioperative metastases formation is largely in agreement with the inhibitory effects of poly(I:C) upon tumour outgrowth in non-surgical models of lung metastases[68]. While polyI:C is ineffective in primates because of inactivation by natural enzymes, other NK stimulators, such as poly-ICLC[69] (stabilized with poly-lysine) or a virus-derived TLR agonist, like the influenza vaccine, could be safely and effectively employed in the perioperative period. Taken together, boosting NK cell activation may counteract the immunosuppressive effects of sepsis and protect against the development of metastatic disease and has potential as a perioperative cancer immunotherapeutic strategy.

## Declarations

### Ethics approval

Animal studies complied with the Canadian Council on Animal Care guidelines and were approved by the University of Ottawa Animal Research Ethics Board.

### Availability of data and materials

Data sharing is not applicable to this article as no datasets were generated or analysed during the current study.

### Author Contributions

**Experimental conception and design** - L-HT, AAA, RS, AA, JZ and CTS.

**Data Acquisition**: L-HT, AAA, RS, AA, JZ and CTS.

**Analysis and interpretation of data (e.g., statistical analysis, biosta-tistics, computational analysis)**: I-HT, AAA, RS, AA, JZ, PS, MAK and RCA.

**Writing, review and revision of manuscript**: L-HT, AAA, RS, PS, MAK and RCA.

**Study Supervision**: L-HT and RCA

## Acknowledgements

Not applicable

